# Microglia protect fungi against copper starvation and promote brain infection

**DOI:** 10.1101/2022.09.07.506901

**Authors:** Sally H. Mohamed, Man Shun Fu, Sofia Hain, Alanoud Alselami, Eliane Vanhoffelen, Yanjian Li, Ebrima Bojang, Robert Lukande, Elizabeth R. Ballou, Robin C. May, Chen Ding, Greetje Vande Velde, Rebecca A. Drummond

**Affiliations:** Institute of Immunology & Immunotherapy, University of Birmingham, Birmingham, UK; Department of Imaging and Pathology, Biomedical MRI/MoSAIC, KU Leuven, Leuven, Belgium; College of Life and Health Sciences, Northeastern University, Shenyang, 110015, Liaoning, China; Department of Pathology, College of Health Sciences, Makerere University, Kampala, Uganda; MRC Centre for Medical Mycology, University of Exeter, UK; Institute of Microbiology & Infection and School of Biosciences, University of Birmingham, Birmingham, UK

## Abstract

Microglia provide protection against a range of brain infections, but how these glial cells respond to fungi is poorly understood. We investigated the role of microglia in the context of cryptococcal meningitis, the most common cause of fungal brain infections in humans. Using a series of transgenic- and chemical-based microglia depletion methods we found that, contrary to their protective role during other infections, microglia supported cryptococcal fungal brain infection. We show that microglia become hosts for intracellular fungal growth and are a site in which the fungus accesses the restricted micronutrient copper. We developed a reporter fungal strain to track copper starvation responses by the fungus and found that yeast were protected from copper starvation within microglia. Lastly, we show that stimulation of microglia with IFNγ causes restriction of phagosomal copper to intracellular fungi. These data provide a mechanistic explanation for why microglia depletion has a therapeutic effect in the context of this life-threatening fungal infection and is one of the few examples of microglia acting to promote infection. Our data demonstrate how tissue-resident phagocytes can support cryptococcal infections by acting as intracellular reservoirs and sites of microbial nutrient acquisition, and how these mechanisms may be blocked by IFNγ immunotherapy.

## Introduction

Tissue-resident macrophages populate every organ in the body and help maintain homeostasis while also acting in an immune surveillance capacity to provide rapid responses to infection or injury. In the central nervous system (CNS), there are several populations of tissue-resident macrophages including meningeal macrophages, perivascular macrophages and microglia (Prinz et al., 2017).

Microglia are the most numerous CNS-resident macrophages. These cells derive from embryonic precursors and maintain their numbers by self-renewal, relying on signals via the CSF1R for survival (Prinz et al., 2017). Microglia are critical for brain development but in adulthood, their main function is immune surveillance and front-line defence against brain infection. The CNS is susceptible to infection with many pathogens. Microglia are ideally situated to respond rapidly to pathogens as they invade the CNS. For example, microglia are required for protective immunity against neurotropic coronaviruses by shaping CD4 T-cell responses within the brain (Wheeler et al., 2018). During infection with the parasite *Toxoplasma gondii*, microglia respond rapidly to produce the alarmin IL-1α and control infection (Batista et al., 2020).

CNS infections with fungi are much rarer than those caused by other microbes, but are difficult to treat and frequently lethal. During brain invasion by *Candida* fungi, microglia depend on CARD9 to activate antifungal immune responses such as the production of neutrophil-attracting chemokines (Drummond et al., 2019). Consequently, human CARD9 deficiency is a primary immunodeficiency disorder that leads to life-threatening CNS *Candida* infections (Drummond et al., 2018).

While many pathogenic fungi are able to infect the human brain, the most common species is *Cryptococcus neoformans*, leading to a disease called cryptococcal meningitis. Cryptococcal meningitis is a leading cause of death in HIV/AIDS patients and is an increasingly observed clinical complication in patients with underlying immune defects caused by iatrogenic interventions (Mohamed et al., 2022). How microglia respond to this fungal infection is unclear. Some studies have observed uptake and survival of the fungus within microglia (Lee et al., 1995), while others have shown robust microglia activation (Neal et al., 2017). Moreover, the microenvironment of the CNS has significant influence over *C. neoformans* morphology and growth responses. For example, *C. neoformans* is severely restricted of the micronutrient copper in the brain, but must deal with toxic copper levels in other organs such as the lung (Sun et al., 2014). Whether tissue-resident macrophages drive these fungal adaptions by regulating access to micronutrients, and the impact of nutritional immunity to organ-specific immune responses, is unknown.

We set out to understand the *in vivo* role of microglia during early infection with *C. neoformans* using a series of microglia depletion strategies. Our findings show that, contrary to other types of infections studied thus far, microglia do not provide protection against this fungal infection. Instead, microglia support cryptococcal brain infection by acting as a growth niche for the fungus and providing access to micronutrients normally restricted within the CNS.

## Results

### Brain macrophages support fungal brain infection

While the role of monocytes and inflammatory macrophages in the control of *C. neoformans* infections has been intensely studied, the role of tissue-resident myeloid cells is less well understood. To definitively determine the contribution of CNS-resident myeloid cells to antifungal immunity, we used several depletion strategies and assessed their effects on control of fungal brain infection. First, we generated *Cx3cr1*-Cre^ER^-iDTR^flox^ animals in which all long-lived CX3CR1^+^ cells in the brain are depleted following treatment with diphtheria toxin (Fig 1A). Microglia highly express CX3CR1, and unlike other CX3CR1+ myeloid cells, do not replenish from the bone marrow. Therefore, we can target expression of the diphtheria toxin receptor (DTR) to only long-lived cells by including a rest period in which peripheral immune cells turn over from the bone marrow (Fig 1A). To examine the role of microglia during experimental *C. neoformans* infection, we used an intravenous infection model where yeast cells directly invade the CNS and bypass pulmonary immunity, allowing us to focus on early host-fungal interactions within the CNS. Using *Cx3cr1*-Cre^ER^-iDTR^flox^ mice, we found that depletion of CX3CR1^hi^CD45^int^ microglia was maintained during fungal infection (Fig 1B). Following *C. neoformans* challenge, we found that the depleted mice had significantly reduced brain fungal burden at 72 hours post-infection but no difference in lung fungal burden (Fig 1C), indicating that microglia depletion may have a protective effect. We also found a significant reduction in brain burdens at day 5 post-infection (Fig S1). To confirm a supportive role for CNS-resident macrophages in *C. neoformans* infection, we employed an independent microglia depletion strategy using the CSF1R inhibitor, PLX5622 (Fig 1E-F), which interrupts the CSF1R-depedent signalling required for microglia survival. We found that PLX5622-treated mice also had reduced fungal brain burdens compared to untreated controls, whereas lung burdens remained unchanged (Fig 1G). Taken together, these data show that depletion of brain-resident macrophages helps to reduce cryptococcal infection specifically in the brain.

**Figure 1:**
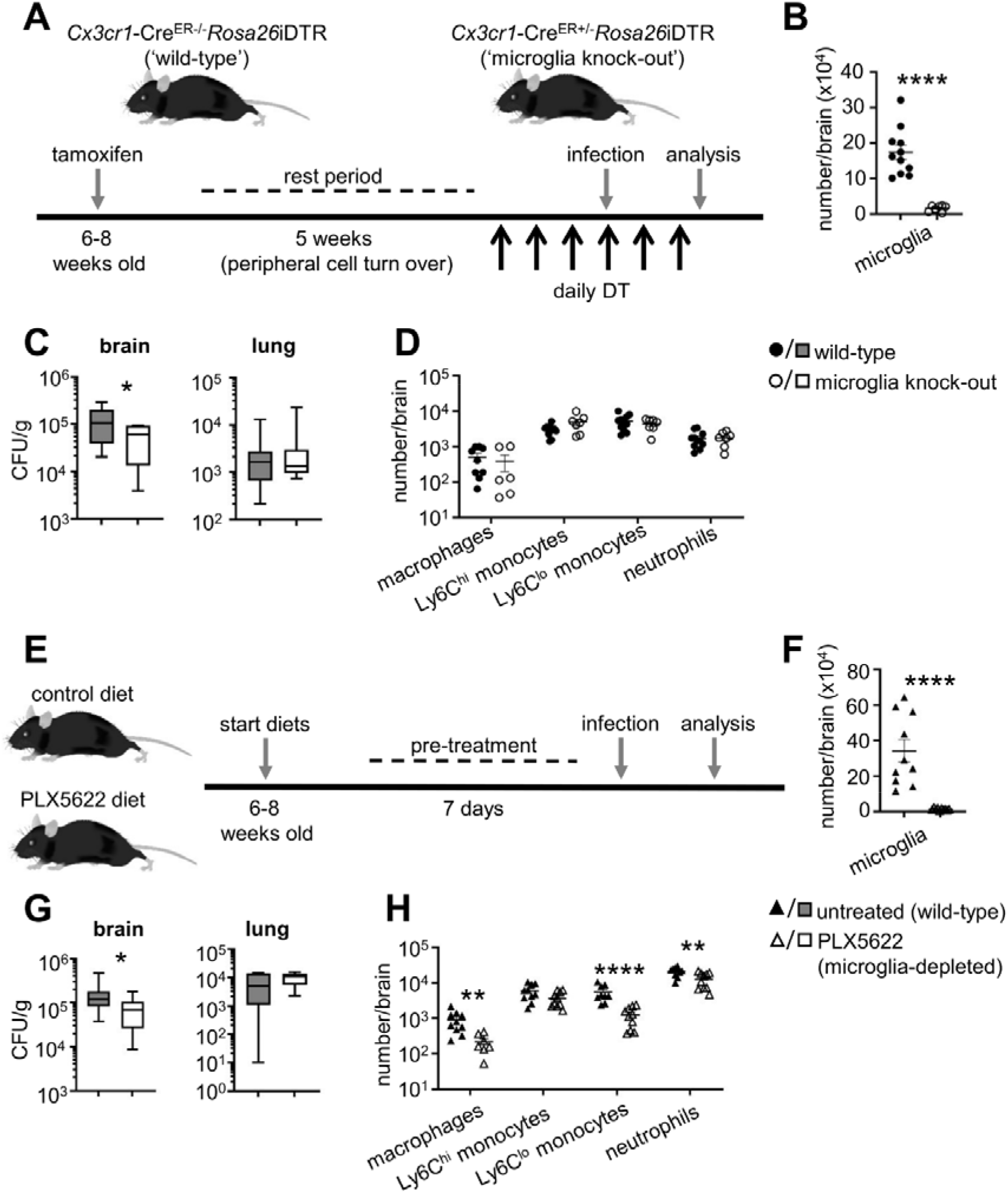
Depletion of CNS-resident macrophages reduces fungal brain burden. (**A**) Schematic of diphtheria toxin-based depletion of brain-resident macrophages. *Cx3cr1*- Cre^ER^ mice were crossed with iDTR mice. Resulting Cre+ (‘wild-type’) and Cre- (‘microglia knock-out’) littermates are treated with tamoxifen to induce Cre expression, and left to rest for 5 weeks to enable turn-over of monocyte-derived macrophages prior to daily treatment with diphtheria toxin to initiate cell depletion before and during intravenous *C. neoformans* H99 infection. (**B**) Total number of microglia in wild-type (n=11) and microglia knock-out (n=7) brains at day 3 post-infection. Data are pooled from 2 independent experiments and analysed by unpaired two-tailed t-test. \*\*\*\**P*<0.0001. (**C**) Fungal burdens in the brain and lung of wild-type (n=11) and microglia knock-out (n=7) mice at day 3 post-infection. Data are pooled from 2 independent experiments and analysed by Mann Whitney U-test. \**P*<0.05. (**D**) Total number of indicated inflammatory cells in the brains of wild-type (n=11) and microglia knock-out (n=6) mice at day 3 post-infection. Data are pooled from 2 independent experiments. (**E**) Schematic of PLX5622 treatment. Wild-type C57BL/6 mice were fed either control diet or PLX5622 diet for 7 days prior to intravenous infection with *C. neoformans* H99. Diets were continued throughout infection. (**F**) Total number of microglia in untreated (n=10) and PLX5622-treated (n=10) brains at day 3 post-infection. Data are pooled from 2 independent experiments and analysed by unpaired two-tailed t-test. \*\*\*\**P*<0.0001. (**G**) Fungal burdens in the brain and lung of untreated (n=14) and PLX5622-treated (n=14) mice at day 3 post-infection. Data are pooled from 3 independent experiments and analysed by Mann Whitney U-test. \**P*<0.05. (**H**) Total number of indicated inflammatory cells in the brains of untreated (n=10) and PLX5622-treated (n=10) mice at day 3 post-infection. Data are pooled from 2 independent experiments.

### Fungal brain infection is primarily supported by microglia

PLX5622 treatment can also deplete other CNS-resident macrophages (Lei et al., 2020; Wheeler et al., 2018) (Fig 1H and Fig S2). Moreover, the *Cx3cr1*-Cre^ER^ line may additionally target non-microglia populations in certain contexts (Masuda et al., 2022; Zhao et al., 2019). Therefore, we sought to generate a mouse model to enable specific depletion of microglia while leaving other CNS-resident and inflammatory macrophages intact, to determine whether the reduction in brain fungal burdens we observed was due to loss of microglia and/or other CNS-resident macrophages.

*Sall1* is a signature microglia gene that is specifically expressed by microglia (Buttgereit et al., 2016). We confirmed this specificity by generating *Sall1*^*CreER*^*R26*^*Ai14*^ transgenic mice, in which only microglia became labelled with dTomato following tamoxifen treatment (Fig 2A-B). Sall1-dependent labelling of microglia was stable during *C. neoformans* infection and remained specific to this cell type (Fig 2C). Therefore, Sall1 is a reliable microglia marker that can be used to specifically target these cells during experimental *C. neoformans* infection. We next crossed *Sall1*^*CreER*^ animals with *Csf1r*^*flox*^ transgenic mice to delete CSF1R specifically in microglia. We found that *Sall1*^*CreER*^*Csf1r*^*flox*^ mice had specific depletion of microglia but normal numbers of other macrophage populations in the CNS after tamoxifen treatment (Fig 2D-E). Importantly, these microglia-specific depleted mice had significantly reduced brain fungal burden and no difference in lung fungal burden (Fig 2F). These data indicate that the reduction in fungal infection observed in the other models is primarily due to the loss of microglia, and that targeted depletion of these cells has a protective effect leading to reduced fungal brain infection.

**Figure 2:**
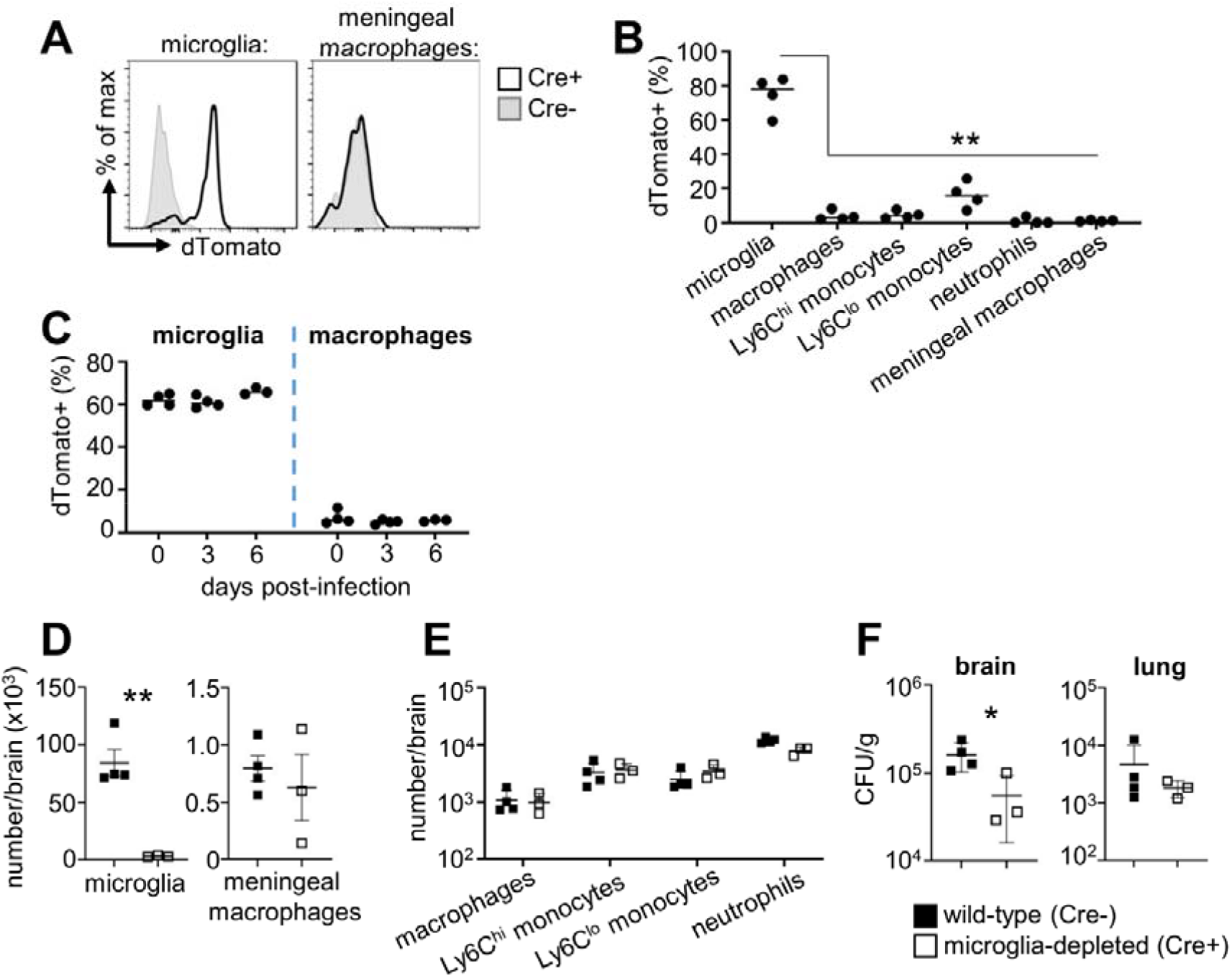
Specific microglia depletion reduces brain fungal burden. (**A**) *Sall1*-Cre^ER^ mice were bred with *Rosa26*^Ai14^ animals and treated with tamoxifen to activate Cre and dTomato expression. Microglia and meningeal macrophage expression of dTomato is shown for representative Cre+ and Cre- littermates. (**B**) Frequency of dTomato expression in the indicated cell populations in the brain in 4 Cre+ littermates. Data are from 1 experiment and analysed by two-way ANOVA. \*\**P*<0.01. (**C**) Frequency of dTomato expression in microglia and brain macrophages in uninfected (n=4) and infected (n=3-4) Cre+ mice. (**D**) Total number of microglia and meningeal macrophages in wild-type (n=4) and microglia-depleted (n=3) brains at day 3 post-infection. Data are representative of 2 independent experiments and analysed by unpaired two-tailed t-test. \*\**P*<0.01. (**E**) Total number of indicated inflammatory cells in the brains of wild-type (n=4) and microglia-depleted (n=3) brains at day 3 post-infection. Data are representative of 2 independent experiments. (**F**) Fungal burdens in the brain and lung of wild-type (n=4) and microglia-depleted (n=3) brains at day 3 post-infection. Data are representative of 2 independent experiments and analysed by unpaired two-tailed t-test. \**P*<0.05

### Microglia depletion does not affect recruitment of inflammatory leukocytes

Microglia can suppress inflammatory responses and activate repair programs to limit tissue damage (Fu et al., 2020; Sariol et al., 2020). We therefore examined whether microglia depletion led to enhanced recruitment of inflammatory myeloid cells to the fungal-infected brain, leading to improved fungal clearance and infection control. We found that microglia depletion did not result in increased recruitment of inflammatory cells to the brain during *C. neoformans* infection in any of the models tested (Fig 1D, 1H, 2E). Taken together, these data show that microglia depletion does not boost inflammatory cell recruitment during experimental *C. neoformans* infection.

### Microglia harbour intracellular fungi

*C. neoformans* can survive and replicate within macrophage phagosomes, aiding its evasion of the immune system and dissemination to the brain within monocytes (Hole and Wormley, 2016). We therefore hypothesised that by depleting microglia we had removed an important growth niche for the fungus within the brain. To examine this possibility, we first characterised the association between fungi and microglia using a green fluorescent reporter *C. neoformans* strain, which allowed us to track fungal uptake by different myeloid cell populations in the brain *in vivo* using flow cytometry (Fig S3). These experiments revealed that ∼5% of microglia became infected with *C. neoformans* (Fig 3A). However, microglia were the most numerous infected cell type within the brain, with significantly greater numbers of infected microglia detected compared to neutrophils, monocytes and inflammatory macrophages (Fig 3B). When examined as a proportion of total infected myeloid cells in the brain, microglia accounted for nearly half of all fungal-infected cells (Fig 3C). Analysis of fungal localisation by histology showed that yeast were growing in the brain parenchyma, in close association or intracellularly in host cells, and within areas of tissue damage with little association with host cells (extracellular growth) (Fig 3D). We also observed similar patterns of growth in the human brain. Pathology analysis of brain tissue from a patient who died from HIV-associated cryptococcal meningitis had areas of extensive tissue damage with large numbers of extracellular yeast, and in other areas, yeast were observed residing within immune cells (Fig 3E). Taken together, this data shows that an intimate relationship exists between tissue-resident microglia and *C. neoformans*.

**Figure 3:**
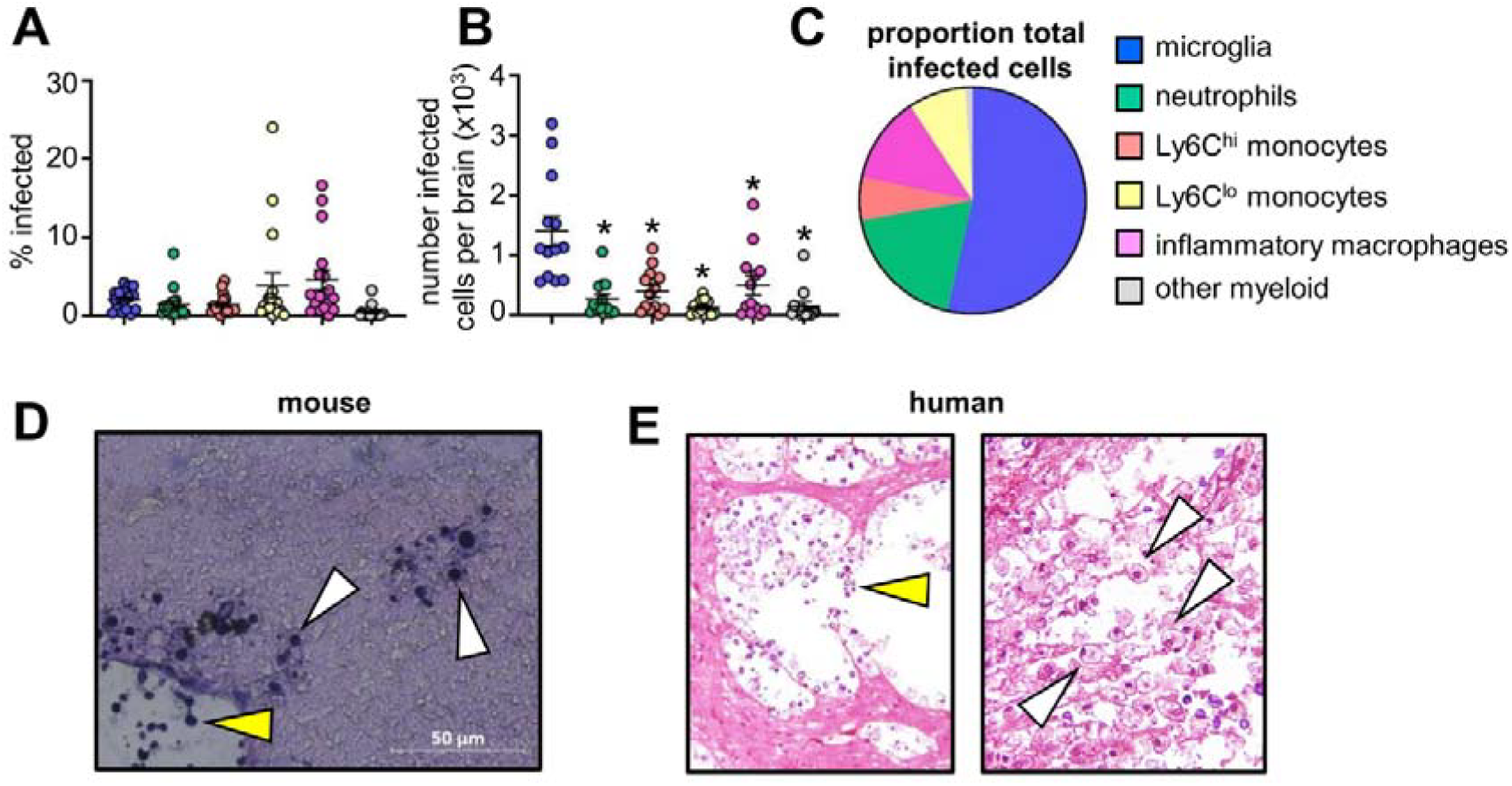
Microglia are hosts to intracellular fungi. (**A**) Frequency of *C. neoformans* infection within indicated cell types in the brain at day 7 post-infection. Data are pooled from 4 independent experiments (n=17 mice). (**B**) Total number of infected cell types in the brain at day 7 post-infection. Data are pooled from 4 independent experiments (n=14 mice) and analysed by one-way ANOVA with Bonferroni correction. \**P*<0.0001. (**C**) Proportion of indicated cell types within total infected cells at day 7 post-infection. An average value of 4 independent experiments for each cell type is shown. See Fig S3 for gating strategies. (**D**) Representative histology of mouse brain at day 6 post- infection, stained with Periodic-Acid Schiff (PAS). White arrows denote intracellular growing yeast, yellow arrows denote extracellular growing yeast. (**E**) Histology of human brain, isolated at autopsy from a patient with HIV-associated cryptococcal meningitis, stained with PAS. White arrows denote intracellular growing yeast, yellow areas denote extracellular growing yeast.

### Intracellular fungal residence within microglia increases brain infection

We next aimed to determine the functional relevance of fungal intracellular survival within microglia. For that, we used the *rdi1*Δ *C. neoformans* mutant, a strain that is unable to survive within macrophages but has normal growth rates in rich media (Price et al., 2008). Histological examination of mouse brains infected with the *rdi1*Δ mutant revealed primarily extracellular yeast in areas of tissue damage at 72h post-infection, whereas wild-type yeast cells were mostly intracellular at this time (Fig 4A). We infected mice that were either microglia-sufficient (untreated) or microglia-depleted (PLX5622) with wild-type or *rdi1*Δ *C. neoformans*, and then quantified brain fungal burden. Compared to wild-type *C. neoformans*, brain fungal burdens were significantly reduced with the *rdi1*Δ strain (Fig 4B). Importantly, while microglia depletion caused a significant reduction in brain infection with wild-type *C. neoformans*, there was no significant difference in brain fungal burdens between microglia-depleted mice and their untreated controls when infected with the *rdi1*Δ mutant (Fig 4B). These data indicate that the reduction in brain fungal burden observed with microglia depletion is dependent on the ability of the fungus to survive intracellularly. We therefore conclude that microglia act as an early growth niche for *C. neoformans* and provide a survival advantage within the brain.

**Figure 4:**
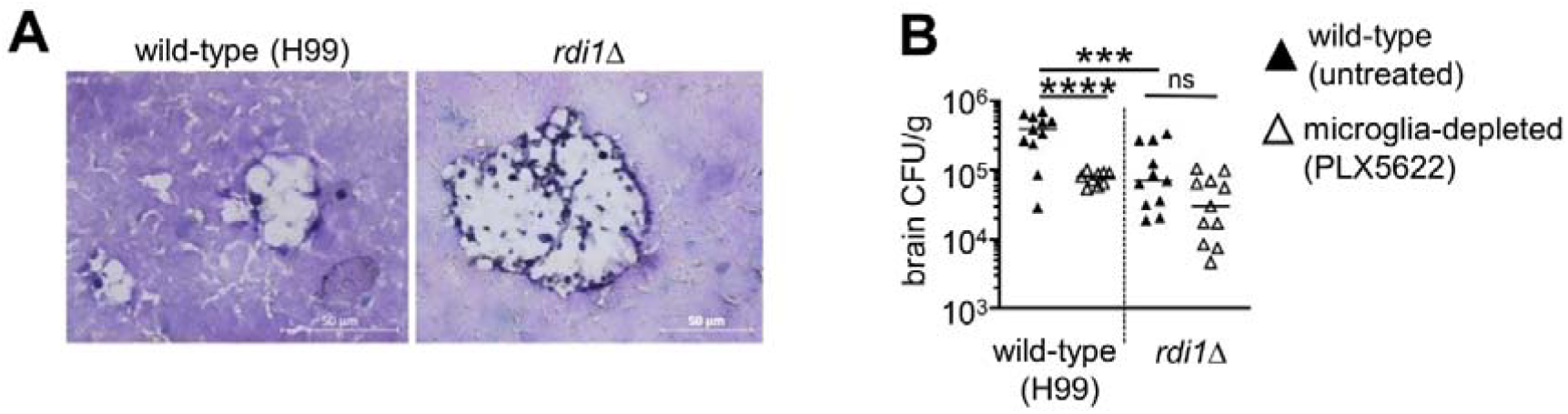
Reduction of fungal brain burden in microglia-depleted mice depends on intracellular fungal growth. (**A**) Representative histology from brain of mice infected with wild-type *C. neoformans* H99 or *rdi1*Δ *C. neoformans* at day 3 post-infection, stained with PAS. (**B**) Brain fungal burdens in untreated or PLX5622-treated mice at day 3 post-infection, infected with either wild-type *C. neoformans* H99 (n=11 wild-type, n=9 microglia-depleted) or *rdi1*Δ *C. neoformans* (n=11 wild-type, n=11 microglia-depleted). Data are pooled from 2 independent experiments and analysed by one-way ANOVA with Bonferroni correction. \*\*\**P*<0.005, \*\*\*\**P*<0.0001. ns = not significant.

### *C. neoformans* is protected from copper starvation within microglia

Macrophages regulate access to micronutrients to either starve or induce toxicity to intracellular pathogens, in a process termed nutritional immunity. Several micronutrients are restricted within the CNS including copper and iron, but whether CNS-resident macrophages participate in this restriction is not well understood. We hypothesised that microglia may be unable to restrict fungal access to micronutrients, thus providing a ‘safe haven’ for *C. neoformans* from starvation conditions found elsewhere in the brain. Indeed, copper is typically found bound within microglia, with extracellular concentrations at 2-3 orders of magnitude lower than intracellular concentrations (Gaier et al., 2013). We therefore tested the cellular localisation of fungal starvation responses in the brain, focusing on copper since *C. neoformans* response to copper starvation in the brain is a driver of virulence (Ding et al., 2013). Under copper-starved conditions, *C. neoformans* upregulates expression of the copper importer *CTR4* (Fig 5A), which is critical for fungal growth and virulence within the CNS (Ding et al., 2013). We compared expression of *CTR4* by yeast cells inside microglia with extracellular yeast *in vivo*, by measuring expression of a GFP transgene under the control of the *CTR4* promotor (*C. neoformans* ^pCTR4^GFP) (Fig 5B). We found that extracellular yeast had upregulated ^pCTR4^GFP expression, in line with previous data that showed significant induction of this gene within the CNS (Sun et al., 2014) (Fig 5B-C). In contrast, we found that yeast associated with microglia had significantly reduced ^pCTR4^GFP expression (Fig 5B-C), indicating that copper concentrations within microglia were adequate for fungal growth. To ensure these results were not an artefact of different fluorescent properties of yeast inside host cells versus free yeast, we normalised ^pCTR4^GFP expression to a constitutively-expressed mCherry housekeeper. Normalised ^pCTR4^GFP expression was similarly reduced in intracellular yeast compared to extracellular yeast (Fig 5D). Taken together, our data show that *C. neoformans* is protected against copper starvation within microglia.

**Figure 5:**
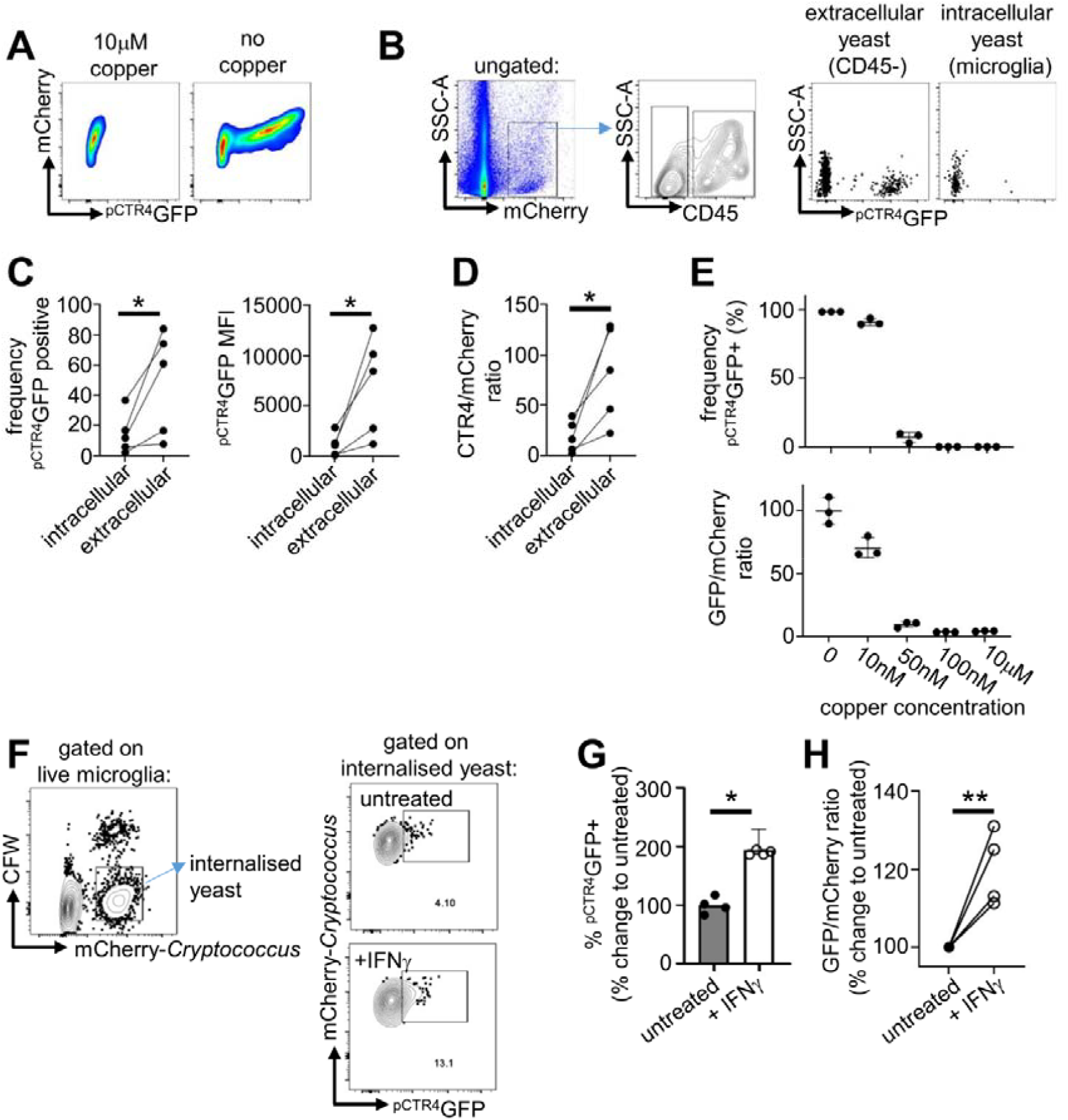
*C. neoformans* is protected from copper starvation when associated with microglia. (**A**) Example flow cytometry plots of ^pCTR4^GFP *C. neoformans* grown in copper-deficient YNB media with and without copper supplementation for 18 hours. mCherry is under the control of the *ACT1* promotor and acts as a housekeeper, GFP expression is controlled by the *CTR4* promotor. (**B**) Gating strategy for analysing *pCTR4-GFP* yeast cells in the brains of mice at day 7 post-infection. Yeast were first gated on using the mCherry marker, then split into extracellular (CD45-) and intracellular (CD45+) groups. Intracellular yeast were further gated to specifically analyse microglia (Ly6G^−^Ly6C^−^CD45^int^CX3CR1^hi^). (**C**) Frequency of GFP+ cells and median fluorescent intensity (MFI) of ^pCTR4^GFP in yeast cells that were extracellular or intracellular within microglia. Each point represents an individual mouse. Data are pooled from two independent experiments and analysed by paired two-tailed t-test. \**P*<0.05. (**D**) Ratio between GFP and mCherry in in yeast cells that were extracellular or intracellular within microglia. Each point represents an individual mouse. Data are pooled from two independent experiments and analysed by paired two-tailed t-test. \**P*<0.05. (**E**) Frequency of GFP+ cells and GFP/mCherry ratio in ^pCTR4^GFP *C. neoformans* grown in copper-deficient YNB media supplemented with indicated concentrations of copper. Yeast cells were analysed by flow cytometry after 18 hours growth. Each point represents a technical replicate. Data are representative of 3 independent experiments. (**F**) Gating strategy used to compare ^pCTR4^GFP expression by intracellular yeast within untreated or IFNγ-treated microglia. Cells were first gated to exclude free yeast, doublets and dead cells. Yeast bound to the surface of microglia but not internalised were removed from analysis using a calcofluor white (CFW) counter stain for the fungal cell wall. (**G**) Frequency of intracellular ^pCTR4^GFP+ yeast in untreated and IFNγ-treated microglia after 2 hours of infection. Bars represent mean data from 4 independent experiments, points show technical replicates from one representative experiment. Data are analysed by unpaired two-tailed t-test. (**H**) Ratio between mCherry and ^pCTR4^GFP expression by intracellular yeast in untreated and IFNγ- treated microglia after 2 hours of infection. Each point represents mean value from 2-3 technical replicates from 4 independent experiments. Data are analysed by unpaired two- tailed t-test. \*\**P*<0.01.

### *C. neoformans* CTR4 expression occurs at <50nM copper

We next used the *pCTR4-GFP C. neoformans* strain to estimate copper concentrations in the CNS. We inoculated copper-deficient growth media with *C. neoformans pCTR4-GFP*, added increasing doses of copper to the media and compared GFP fluorescence after 24 hours growth by flow cytometry. At 10μM copper, there was no detection of ^pCTR4^GFP expression and yeast cells remained negative for GFP expression down to 100nM copper (Fig 5E). We began to detect expression of ^pCTR4^GFP at 50nM copper, where ∼5% yeast cells were GFP+ (Fig 5E). However, at 10nM copper this rose significantly to ∼90% GFP+, similar to the copper starved condition (Fig 5E). These data indicate that *CTR4* expression switches on at copper concentrations <50nM and does not exhibit a graded expression across a range of copper concentrations. Based on these data, we estimate that copper concentrations in the extracellular compartment of the *C. neoformans*-infected brain are between 10-50nM, whereas copper concentrations within *C. neoformans*-infected microglia are 50-100nM or higher.

### IFNγ induces copper restriction in fungal-infected microglia

IFNγ is a pro-inflammatory cytokine that mediates crosstalk between CD4 T-cells and macrophages to initiate protective immunity against *C. neoformans* infection (Leopold Wager et al., 2015; Williamson et al., 2017). IFNγ has also been shown to modulate copper concentrations within macrophage phagosomes inducing either copper deprivation (Shen et al., 2018) or importing excess copper to induce toxicity (Djoko et al., 2015). Whether IFNγ modulates copper levels within microglia and if this contributes towards the protective effects observed in anti-cryptococcal immunity has not been explored. We therefore sought to understand whether IFNγ influenced intracellular copper concentrations in microglia and impacted starvation responses by intracellular fungi. We infected the murine microglia cell line BV-2 with *C. neoformans pCTR4-GFP*, comparing ^pCTR4^GFP expression by yeast internalised within untreated microglia, or microglia pre-treated with IFNγ (Fig 5F). We found that yeast internalised by IFNγ-treated microglia had significantly higher expression of ^pCTR4^GFP than yeast internalised by untreated microglia (Fig 5G,H). These data suggest that IFNγ induces copper restriction within microglia phagosomes resulting in an enhanced fungal copper starvation response.

## Discussion

Microglia have been shown to play a central role in the protection against CNS infections, including *Candida albicans* (Drummond et al., 2019), *Toxoplasma gondii* (Batista et al., 2020) and viruses (Wheeler et al., 2018). Here, we show that microglia have the opposite effect in cryptococcal meningitis and support early *C. neoformans* growth in the brain by subverting nutritional immunity.

Microglia can rapidly respond to microbial invasion of the CNS. For example, following *C. albicans* fungal brain infection, microglia produced IL-1β and CXCL1 within 24 hours of infection, which elicited a significant influx of neutrophils to the brain to enable clearance of the infection (Drummond et al., 2019). During infection with the parasite *T. gondii*, microglia produced IL-1α which was required for the protective recruitment of inflammatory T-cells (Batista et al., 2020). In our work, we found no consistent evidence for a role of microglia in recruiting inflammatory myeloid cells to the CNS during *C. neoformans* infection. This lack of a role for microglia in early protective anti-cryptococcal immunity may be linked with poor induction of pro-inflammatory immune signalling in response to *C. neoformans*. Many fungi engage CARD9-coupled C-type lectin receptors on host immune cells, leading to a pro-inflammatory signalling cascade and leukocyte activation (Drummond et al., 2018). However, *C. neoformans* secretes a mannose-rich capsule which shields the fungal cell wall from recognition by CARD9-coupled receptors, such as Dectin-1 (Walsh et al., 2017). Indeed, in contrast to other fungal infections, patients with deficiencies in CARD9 and related molecules do not appear to be more susceptible to cryptococcal meningitis (Drummond et al., 2018), indicating that these pathways are redundant for protection against *C. neoformans* infection. The innate receptors mediating activation of brain-resident macrophages during *C. neoformans* infection remains to be determined, but this process likely depends on protective crosstalk with IFNγ-producing CD4 T-cells. In a model of chronic *C. neoformans* brain infection, microglia were activated after significant infiltration of pathologic IFNγ-producing CD4 T-cells which subsequently drove neuroinflammation and death (Neal et al., 2017). In our study, we have used an acute infection model and examined early time points (<1 week) prior to significant T-cell infiltration. Our data show that microglia are an important reservoir for early infection, and that stimulation with IFNγ may help limit intracellular infection of these cells by restriction of phagosomal copper.

In addition to hosting intracellular fungi, microglia may also support infection by enabling fungal invasion across the blood-brain-barrier (BBB). *C. neoformans* is thought to invade the CNS using a variety of routes, including transcellular invasion of the BBB as free yeast or within host monocytes (Santiago-Tirado et al., 2017). Whether microglia are involved with either of these processes is unclear. A small subset of microglia, called capillary-associated microglia (CAMs), have been shown to associate with blood vessels in the brain and are involved in maintaining integrity via purine signalling (Bisht et al., 2021). During sterile neuroinflammation, microglia migrate to cerebral blood vessels to regulate BBB integrity but can contribute to barrier breakdown when inflammation is inappropriately sustained (Haruwaka et al., 2019; Yu et al.). Loss of BBB integrity has been previously observed during cryptococcal meningitis, which is associated with fungal proliferation within cerebral blood vessels (Gibson et al., 2022). The role of microglia in BBB integrity during cryptococcal meningitis is still unresolved, and we cannot rule out that the reduction in brain fungal burdens observed in our microglia-depleted mice is partially due to effects on BBB integrity caused by depletion of CAMs and/or interruption of inflammatory microglia migration to the blood vessels. Future studies should examine microglia subsets and migration patterns during fungal infection to identify the different ways in which these cells contribute towards pathology.

Some of the microglia depletion methods used in our study, including the CSF1R inhibitor PLX5622 (Kerkhofs et al., 2020; Spiteri et al., 2022), are reported to have off target effects on border macrophages within the CNS. The role of non-microglia CNS border macrophages in the control and dissemination of *C. neoformans* is not fully understood, but they have been shown to become heavily infected with the fungus (Kaufman-Francis et al., 2018). Some of the depletion models used in our study may have interrupted fungal invasion tactics that involve these non-microglia populations, contributing to the reduction in fungal brain burdens we observed. To test this possibility, we generated *Sall1*^*CreER*^*Csf1r*^*flox*^ animals, which had specific depletion of microglia and reduced fungal brain burdens, indicating that microglia are the main macrophages within the CNS that promote fungal brain infection. Further study is required to specifically examine the function of rarer border macrophages in cryptococcal meningitis and determine whether they contribute towards CNS dissemination and/or prevent fungal invasion across the BBB.

When we explored how microglia supported *C. neoformans* infection, we discovered that the fungus was protected from copper starvation within these cells. Copper is an essential heavy metal critical for many biochemical processes (Desai and Kaler, 2008). In the CNS, intracellular copper levels are up to 1000-fold higher than extracellular levels (Gaier et al., 2013). This is likely because copper build-up within the CNS is toxic, leading to neuroinflammation of which inappropriate microglia activation is a hallmark (Desai and Kaler, 2008). For example, high copper concentrations and dysregulated microglia expression of copper transporters has been measured in Alzheimer’s plaques (Lovell et al., 1998; Zheng et al., 2010). Due to the restricted copper availability within the CNS, *C. neoformans* significantly upregulates copper importing proteins Ctr1 and Ctr4 within the brain (Ding et al., 2013; Sun et al., 2014). Indeed, *C. neoformans* mutants deficient in *CTR4* have attenuated growth and virulence specifically within the CNS (Ding et al., 2013; Sun et al., 2014).

We tracked *C. neoformans* copper sensing using a *pCTR4-GFP* reporter strain. We found that *C. neoformans* highly upregulated CTR4 in the brain following intravenous infection, confirming previous work that showed *C. neoformans* is starved of copper in this tissue (Ding et al., 2013; Garcia-Santamarina et al., 2020; Sun et al., 2014). However, we found that CTR4 induction was significantly lower in yeast that were associated with microglia. These data indicate that copper concentrations in microglia were adequate for fungal growth. Similar observations have been made with the intracellular replicating fungus *Histoplasma capsulatum*, which uses the copper importer Ctr3 to protect against copper starvation (Shen et al., 2018). Infection of bone-marrow derived macrophages *in vitro* showed that *H. capsulatum* does not upregulate *CTR3* in the phagosome of resting macrophages and can acquire copper from this site, enabling its intracellular infection program (Shen et al., 2018). IFNγ stimulation of macrophages caused restriction of phagosomal copper concentrations, activating a copper-starvation response in *H. capsulatum* marked by upregulation of the copper importer *CTR3* (Shen et al., 2018). In this case, IFNγ may partly mediate protective immunity by limiting intracellular copper availability. Our data point to a similar mechanism occurring in IFNγ-stimulated microglia, helping to limit fungal intracellular growth in the brain. Adjunctive therapy with recombinant IFNγ in patients with HIV-associated cryptococcal meningitis decreased fungal burden in the CSF, a marker correlated with survival (Jarvis et al., 2012). The specific protective mechanisms of IFNγ in anti-cryptococcal protection in the CNS are not fully resolved. Our data indicate that IFNγ may exert protective effects directly on microglia, at least partially, by limiting fungal access to key nutrients. Future studies will need to examine other nutrients that the fungus may access within microglia, such as iron and lipids, which have both been shown to be acquired by *C. neoformans* within other macrophage types (Kronstad et al., 2013; Nolan et al., 2017).

In summary, our data highlight microglia as an important reservoir for intracellular fungal infection, acting to shield the fungus from the copper-deficient environment within the CNS. We also show that targeting microglia with IFNγ may be beneficial by boosting nutritional immunity in these tissue-resident macrophages to limit their supportive role in brain infection. This is one of the first examples of microglia acting to promote CNS infection, and delivers new insights into critical pathways underlying host-fungal interactions in the brain during cryptococcal meningitis.

## Methods

### Mice

8-12 week old mice (males and females) were housed in individually ventilated cages under specific pathogen free conditions at the Biomedical Services Unit at the University of Birmingham, and had access to standard chow and drinking water *ad libitum*. Animal studies were approved by the Animal Welfare and Ethical Review Board and UK Home Office under Project Licence PBE275C33. Wild-type refers to C57BL/6 (Charles River) or the corresponding littermates of genetically-modified lines; *Cx3cr1-*Cre^ER^*Rosa26*iDTR, *Sall1*- Cre^ER^*Rosa26*^Ai14^, and *Sall1*-Cre^ER^*Csf1r*^flox^. *Cx3cr1*-Cre^ER^, *Rosa26*iDTR, *Rosa26*^Ai14^ and *Csf1r*^flox^ mice were originally purchased from Jackson and colonies bred and maintained at the University of Birmingham. *Sall1*-Cre^ER^ mice were a kind gift from Dr Melanie Greter (University of Zurich). Mice were euthanised by cervical dislocation at indicated analysis time-points, or when humane endpoints (e.g. 20% weight loss, hypothermia, meningitis) had been reached, whichever occurred earlier.

### Tamoxifen and diphtheria toxin treatments

Tamoxifen was dissolved in 100% ethanol (1g/mL) and diluted in corn oil (Sigma) to 100mg/mL. Mice received two doses of tamoxifen by oral gavage (10mg in 100μL) 48 hours apart, then were left to rest for at least 1 week (experiments using Sall1-Cre^ER^ crossed strains). In some experiments, mice were left to rest for 5-6 weeks prior to dosing with diphtheria toxin (experiments using Cx3cr1-Cre^ER^ crossed strains). Diphtheria toxin (Sigma) was injected intraperitoneally (30ng per gram body weight) daily for 3 days prior to infection, on the day of infection, and for 2 days after.

### PLX5622 treatment

PLX5622 (Plexxikon Inc. Berkley, CA) was formulated in AIN-76A rodent chow (Research Diets) at a concentration of 1200 mg/kg. Mice were provided with PLX6522 diet or AIN-76A control diet *ad libitum* for 1 week prior to infection, and continued throughout the infection study period.

### *C. neoformans* growth and mouse infections

*C. neoformans* strains used in this study were H99, KN99α-GFP(Voelz et al., 2010), *rdi1*Δ(Price et al., 2008), and *pCTR4-GFP* (this study). Yeast was routinely grown in YPD broth (2% peptone [Fisher Scientific], 2% glucose [Fisher Scientific], and 1% yeast extract [Sigma]) at 30 °C for 24 hours at 200rpm. In some experiments, *C. neoformans pCTR4-GFP* were grown in copper-deficient YNB media (Formedium), supplemented with 2% glucose and copper sulphate (Sigma; see figure legends for specific concentrations). For infections, yeast cells were washed twice in sterile PBS, counted using haemocytometer, and 2×10^4^ yeast injected intravenously into the lateral tail vein. For analysis of brain and lung fungal burdens, animals were euthanized and organs weighed, homogenized in PBS, and serially diluted before plating onto YPD agar supplemented with Penicillin/Streptomycin (Invitrogen). Colonies were counted after incubation at 37**°**C for 48 hours.

### Generation of *pCTR4-GFP C. neoformans*

The *pCTR4-GFP-mCherry* strain was generated by transforming the integrative *pCTR4-GFP* plasmid (Sun et al., 2014), which harbours GFP encoding gene driven by copper deficiency inducible promotor (*CTR4*), into KN99α-mCherry. The mCherry strain was a gift from Dr. Tongbao Liu (Southwest University, China), mCherry expression is controlled by the *ACT1* promotor. Transformations of *C. neoformans* were completed using biolistic transformation, as previously described(Toffaletti et al., 1993). The resulting *pCTR4-GFP-mCherry* strain was validated through growth on copper-sufficient and copper-deficient media to check correct functioning of the ^pCTR4^GFP construct.

### Analysis of Brain Leukocytes by FACS

Leukocytes were isolated from brain using previously described methods. Briefly, brains were aseptically removed and stored in ice-cold FACS buffer (PBS + 0.5% BSA + 0.01% sodium azide) prior to smashing into a paste using a syringe plunger. The suspension was resuspended in 10mL 30% Percoll (GE Healthcare), and underlaid with 1.5mL of 70% Percoll. Gradients were centrifuged at 2450 rpm for 30 min at 4 °C with the brake off. Leukocytes at the interphase were collected and washed in FACS buffer prior to labelling with fluorophore-conjugated antibodies and flow cytometry analysis.

### Flow Cytometry

Isolated leukocytes were resuspended in PBS and stained with Live/Dead stain (Invitrogen) on ice as per manufacturer’s instructions. Fc receptors were blocked with anti-CD16/32 and staining with fluorochrome-labelled antibodies was performed on ice. Labelled samples were acquired immediately or fixed in 2% paraformaldehyde prior to acquisition. Anti-mouse antibodies used in this study were: CD45 (30-F11), CD11b (ICCRF44), CX3CR1 (SA011F11), MHC Class II (M5/114.15.2), F480 (BM8), Ly6G (1A8), Ly6C (HK1.4), all from Biolegend. Samples were acquired on a BD LSR Fortessa equipped with BD FACSDiva software. Analysis was performed using FlowJo (v10.6.1, TreeStar).

### Tissue Culture

BV2 cells (a kind gift from Dr Michail Lionakis, NIH) were routinely maintained at 37 °C, 5% CO_2_ in RPMI (supplemented with GlutaMax and HEPES, Gibco), further supplemented with 10% heat-inactivated foetal bovine serum (Gibco) and 1% Penicillin/Streptomycin (Invitrogen), and were split every 2-3 days after reaching 80-90% confluence. For experiments, BV2 cells were lifted using Trypsin-EDTA (Sigma) and a cell scraper, counted using Trypan blue exclusion, and seeded into 12 well plates, at 2×10^5^ cells per well in 1mL media. In some wells, media was further supplemented with 100ng/mL recombinant mouse IFNγ (Biolegend). After 24 hours, BV2 cells were infected with 1×10^6^ *C. neoformans* CTR4^pGFP^ yeast that had been pre-opsonised with 10μg/mL anti-glucuronoxylomannan (GXM) antibody (18B7; Millipore) for 15 minutes at room temperature. After 2 hours, plates were placed on ice, media removed and replaced with 1mL ice-cold 2mM EDTA in 1xPBS. BV2 cells were lifted by gentle pipetting on ice and transferred to FACS tubes prior to staining with fluorophore-conjugated antibodies and 5μg/mL calcofluor white (Sigma), then acquired immediately on a BDFortessa as above.

### Histopathology

Mouse brains were removed from infected mice at indicated time points and either fixed in 10% formalin for 24 hours before embedding in paraffin wax, or frozen in OCT before sectioning. Tissue sections were stained with periodic Acid-Schiff (PAS) and hematoxylin and eosin (H&E). Imaging of tissue sections were captured using a Zeiss Axio Slide Scanner and data analysed using Zen Lite Blue (version 1.1.2.0).

### Statistics

Statistical analyses were performed using GraphPad Prism 9.0 software. Details of individual tests are included in the figure legends. In general, data were tested for normal distribution by Kolmogorov-Smirnov normality test and analyzed accordingly by unpaired two-tailed *t*-test or Mann Whitney *U*-test. In cases where multiple data sets were analyzed, two-way ANOVA was used with Bonferroni correction. In all cases, *P* values <0.05 were considered significant.

## Acknowledgements

We would like to thank the technical staff at the Biomedical Services Unit (Birmingham) for their care and help with animal husbandry. We thank Dr Guillaume Desanti and support staff at the Flow Cytometry Unit at the University of Birmingham for their support with sorting and flow cytometry experiments. This work was funded by the Academy of Medical Sciences (SBF004_1008, awarded to RAD), Medical Research Council (MR/S024611, awarded to RAD), Research Foundation Flanders (PhD studentship awarded to EV, 1SF2222N) and the National Natural Science Foundation of China (31870140, awarded to CD).

---

**Figure S1.**
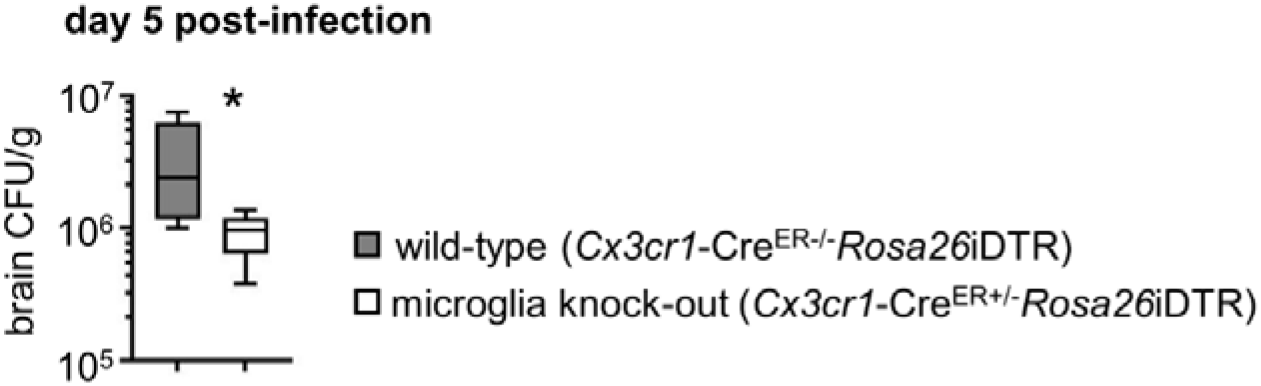
Brain fungal burdens at day 5 post-infection in wild-type (n=7) and microglia-deficient (n=5) mice (see Figure 1 legend for details). Data is pooled from 2 independent experiments and analysed by Mann Whitney U-test. \**P*<0.05

**Figure S2.**
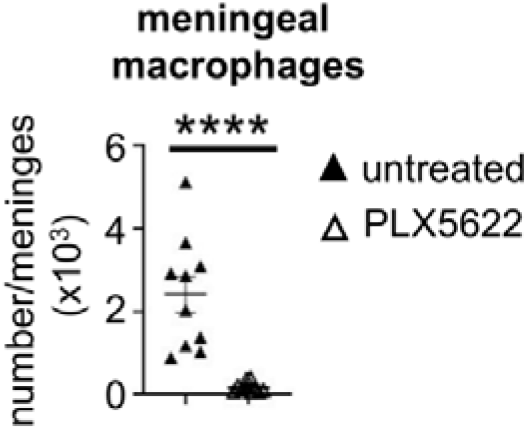
Number of meningeal macrophages in untreated (n=10) and PLX5622- treated (n=10) C57BL/6 mice at day 3 post-infection. Data is pooled from 2 independent experiments and analysed by unpaired two-tailed t-test. \*\*\*\**P*<0.0001

**Figure S3.**
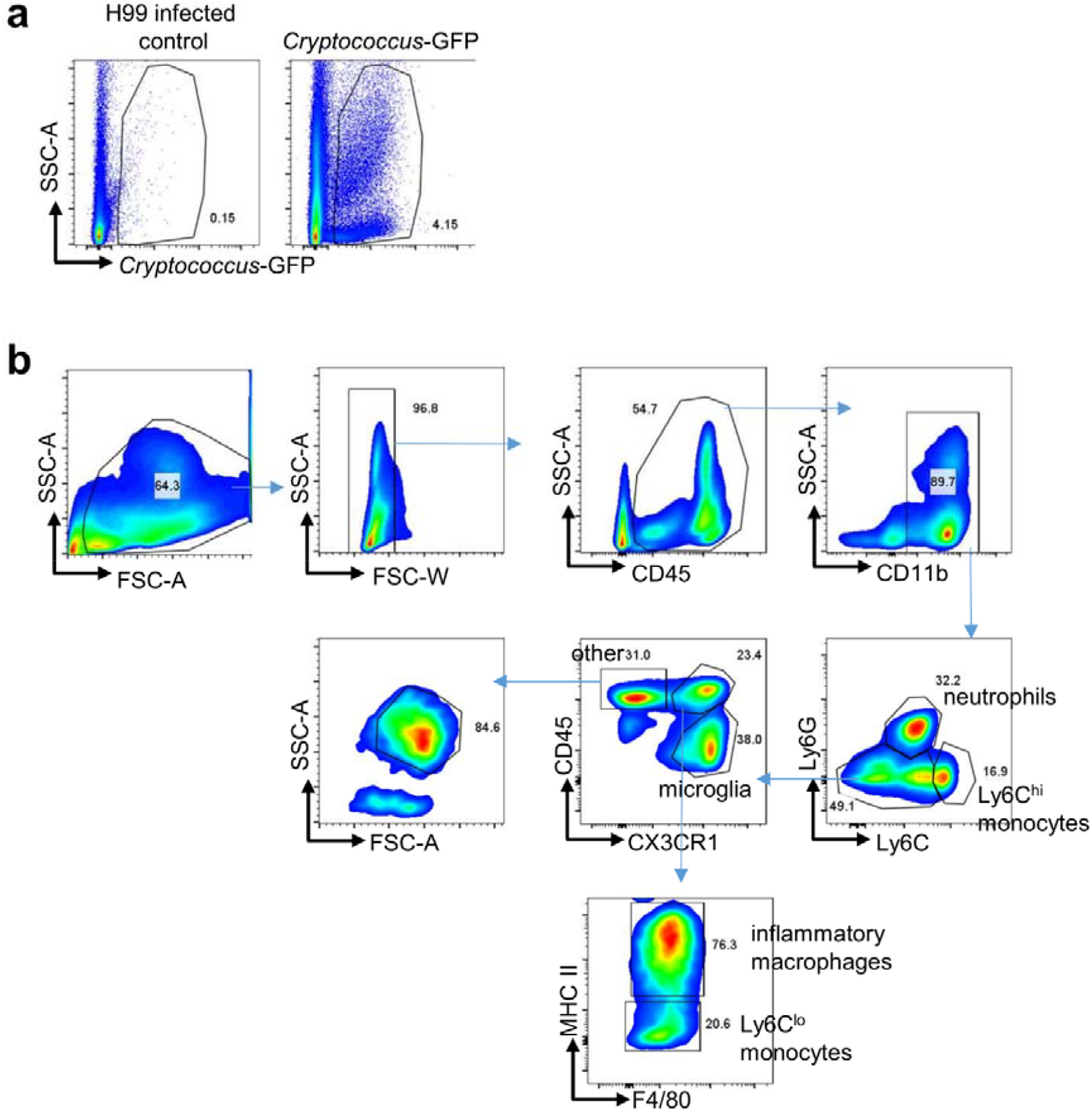
Gating strategies used to analyse (**a**) yeast population or (**b**) host leukocytes in the infected mouse brain. In (a), plots are ungated and show examples from mice infected with non-fluorescent control H99 (as a gating control) and GFP- expressing *C. neoformans*. In (b), the main immune cell populations routinely gated on in our experiments are shown. Within the ‘other’ gate, we found that most cells are SSC^high^ and are likely eosinophils.

## Notes

### Competing Interest Statement

The authors have declared no competing interest.

